# Effects of pasteurization on osteopontin levels in human breastmilk and pasteurized breastmilk products

**DOI:** 10.1101/2023.02.24.529945

**Authors:** Kathleen G. McClanahan, Jeff Reese, Jörn-Hendrik Weitkamp, Danyvid Olivares-Villagómez

## Abstract

**Background:** Osteopontin (OPN) is an important breastmilk protein involved in infant intestinal, immunological, and brain development. However, little is known about how common milk pasteurization and storage techniques affect this important bioactive protein.

**Methods:** Human milk osteopontin concentration was measured in single-donor fresh or frozen breastmilk, pooled Holder-pasteurized donor breastmilk, and a shelf-stable (retort pasteurized) breastmilk product by ELISA. Breastmilk samples were pasteurized and/or frozen before measuring osteopontin concentrations.

**Results:** Holder pasteurization of breastmilk resulted in an ∼50% decrease in osteopontin levels within single-donor samples, whereas pooled donor breastmilk had comparable osteopontin levels to non-pasteurized single-donor samples. Breastmilk from mothers of preterm infants trended toward higher osteopontin concentration than mothers of term infants; however, samples from preterm mothers experienced greater osteopontin degradation upon pasteurization. Finally, freezing breastmilk prior to Holder pasteurization resulted in less osteopontin degradation than Holder pasteurization prior to freezing.

**Conclusion:** Commonly used breastmilk pasteurization and storage techniques, including freezing, Holder and retort pasteurization, decrease the levels of the bioactive protein osteopontin in human breastmilk.

**Impact:**

- Pasteurization of human breastmilk significantly decreases the levels of the bioactive protein osteopontin
- Use of both pasteurization and freezing techniques for breastmilk preservation results in greater loss of osteopontin
- This study presents for the first time an analysis of osteopontin levels in single-donor pasteurized milk samples

## Introduction

Preterm birth is the leading cause of neonatal morbidity and mortality worldwide.^1^ In the United States, about 10% of infants are born prematurely (<37 weeks gestation)^2^ and have increased susceptibility to infection,^3^ as well as complications including sepsis, retinopathy of prematurity, and necrotizing enterocolitis (NEC). It is well established that providing breastmilk is a preventative measure against the development of NEC and other complications in the premature infant.^4,5,6,7^

The gold standard for premature infant nutrition is Mother’s own Milk (MoM), whose components fluctuate to meet the nutritional and immunological needs of the infant. However, in many instances MoM is not available, and pooled donor breastmilk is commonly used as a substitute. Pooled donor breastmilk is a combination of milk from multiple donors that has been Holder pasteurized (30 minutes at 62.5°C), and has been shown to improve outcomes in premature infants compared to infant formula.^8^ Another source of nutrition is shelf-stable donor breastmilk products, which are usually retort pasteurized, a harsher method of pasteurization (115-145°C under high pressure for several minutes).

Breastmilk contains many bioactive components including oligosaccharides, IgA, and proteins such as lactoferrin and lysozyme that play an important role in supporting infant immunity while the neonatal immune system begins to develop.^9^ However, pasteurization processes, including Holder, retort, and others (e.g. high-temperature short-time pasteurization and high pressure processing), subject breastmilk to conditions that have been shown to significantly affect the bioactivity of these components. For example, Holder pasteurization decreases the activity of IgA, lactoferrin, and lysozyme.^10,11,12^ In addition, retort pasteurization has been shown to further decrease the bioactivity of these important milk components.^13,14,15^ Although pasteurization is important to ensure breastmilk safety, discussion of the impacts of breastmilk sterilization has been on the rise.^16,17,18^

Recently, attention has been drawn toward another bioactive milk component: osteopontin, which has been shown to play a role in infant immune and intestinal development.^19,20,21^ Osteopontin is a highly phosphorylated glycoprotein that plays pleiotropic roles, including in cell adhesion,^22^ immune cell modulation,^23^ and prevention of calcification.^24^ Osteopontin is expressed by many cell types, and is present in bodily fluids, including blood, urine, and milk. The concentration of osteopontin in human milk varies depending on the milk stage (18 to 322 mg/L), with higher levels in colostrum and transitional milk, and lower levels in mature milk.^25^ The concentration of breastmilk osteopontin fluctuates considerably among women; in a study performed across three different countries, Danish women presented the lowest levels of breastmilk-derived osteopontin (99.7 mg/L), while Japanese and Chinese women of similar socioeconomic status presented 185 and 266 mg/L, respectively.^26^ Milk-derived osteopontin has been shown to move intact through the gastrointestinal tract, and enter the infant’s circulation, suggesting a systemic role for this protein. Previous studies in human infants have demonstrated that milk osteopontin plays a role in infant immune development, ^20,21,25^ with further animal studies implicating additional roles in brain and intestinal development. ^19,27^

Due to the potential importance of osteopontin, it is critical to determine how pasteurization affects its levels in breastmilk. It has been noted that Holder and retort pasteurization affect the levels of osteopontin in pooled donor breastmilk.^28^ However, this report analyzed a single sample of previously-frozen pooled breastmilk, which may not fully represent the variance in other breastmilk sources. Thus, in this report, we investigated the levels of osteopontin in human breastmilk from different sources, and how pasteurization and freezing impact the concentration of this important bioactive component.

## Methods

### Sample acquisition

Deidentified human breastmilk samples were acquired from subjects at Vanderbilt University Medical Center in Nashville, Tennessee, U.S.A. This project was approved by the Institutional Review Board at Vanderbilt University and all participants gave informed consent for their samples to be used for research. Samples were divided into groups based on infant gestational age at birth: preterm (<37 weeks gestation) and term (>37 weeks gestation). Single-donor samples were stored at −80°C until use. Samples of pooled donor breastmilk and retort pasteurized shelf-stable human breastmilk (Ni-Q HDM Plus, Ni-Q, Wilsonville, OR) were acquired from the milk feeding preparation room at Monroe Carrel Jr. Children’s Hospital at Vanderbilt in Nashville, Tennessee and stored at 4°C or room temperature, respectively, until use. Frozen individual samples of donor breastmilk from mothers of preterm and term infants were thawed and subjected to Holder pasteurization. We measured osteopontin levels before and after freezing and pasteurization. Fresh breastmilk samples were stored at 4°C until use.

### Milk Processing

Single-donor frozen milk samples were thawed in a room-temperature water bath. Samples were divided into two 5mL aliquots – one of which was subjected to Holder pasteurization in a 62.5°C water bath for 30 minutes, and the other was left at 4°C for 30 minutes. Fresh donor milk was divided into 400 µl aliquots before being subjected to Holder pasteurization and/or freezing. Frozen samples were thawed in a room-temperature water bath prior to analysis.

### Determination of Protein Concentration

The concentration of osteopontin in human milk samples was determined in triplicate using the Human Osteopontin ELISA Kit (R&D Systems, Minneapolis, MN). Samples were diluted 1:100,000 in the provided buffer and analyzed following the manufacturer’s instructions.

### Statistical Analysis

Data were analyzed using Prism 9 (GraphPad, San Diego, CA) to determine significant (p < 0.05) differences between groups. Comparisons between two groups were made by Student’s t-test and comparisons between multiple groups were made using one-way analysis of variance (ANOVA) with post hoc Tukey’s test. Data are shown as average ± standard error of the mean. Comparisons between pre- and post-pasteurized samples were made using Student’s paired t-test.

## Results

We investigated how Holder pasteurization affects the levels of osteopontin in human breastmilk. For this purpose, we obtained frozen breastmilk samples from different donors and observed that the levels of osteopontin were, as expected, variable among samples (Figure 1). While the average was ∼50 µg/ml, some samples contained little to no osteopontin, and others had levels exceeding 100 µg/ml. When the samples were Holder pasteurized, the average osteopontin concentration dropped by 50% to 25 µg/ml, which is comparable to the decrease in other bioactive proteins reported in the literature.^12^ Of note, many samples had no detectable osteopontin after treatment. On the other hand, pooled pasteurized breastmilk had a similar average osteopontin level as single-donor breastmilk without pasteurization. Interestingly, none of the pooled samples analyzed had osteopontin concentration below 25 µg/ml. Shelf-stable breastmilk, which undergoes the more stringent retort pasteurization, presented very low levels of osteopontin.

**Figure 1:**
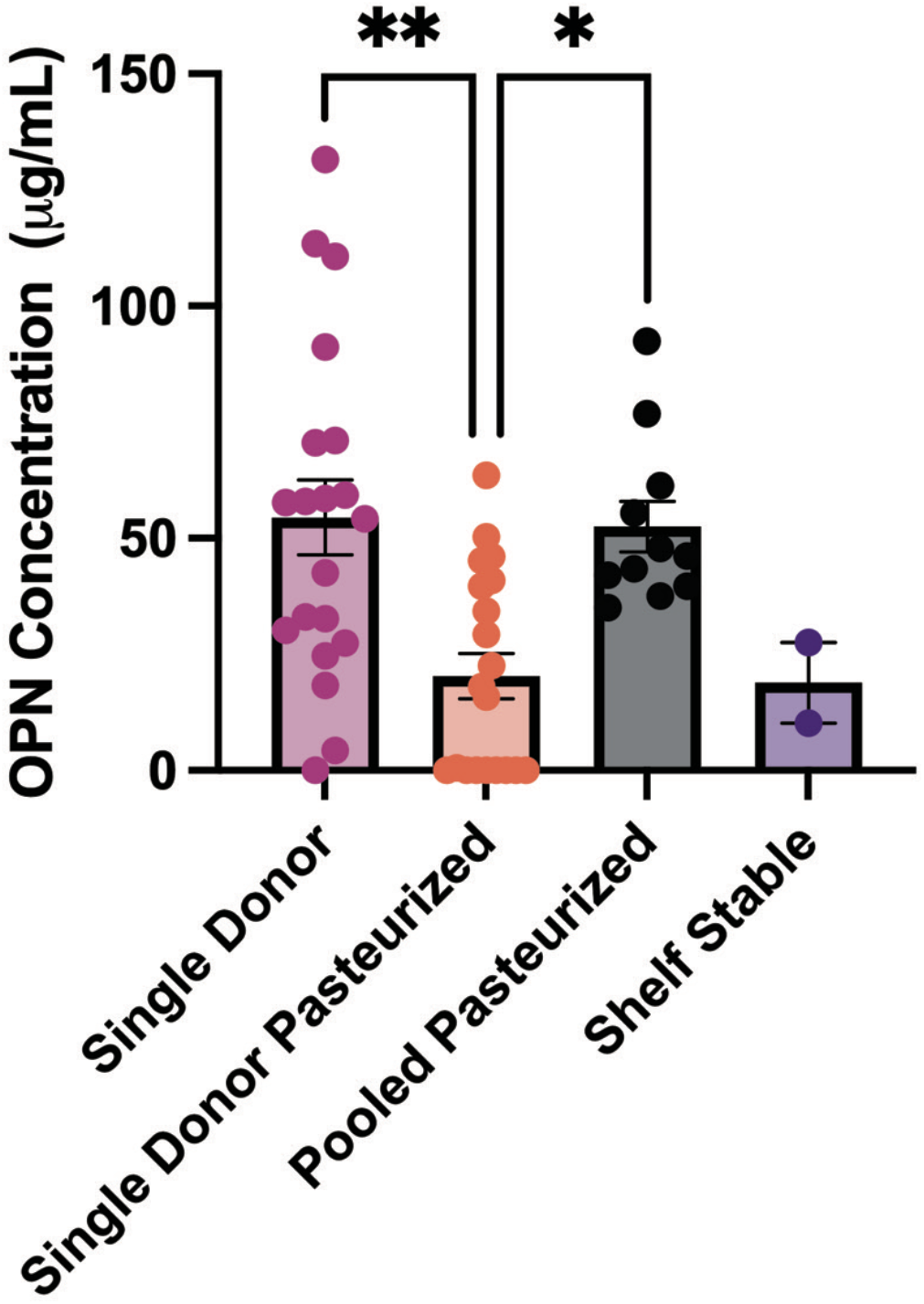
Osteopontin levels in the indicated breastmilk products. Human milk osteopontin concentrations were measured using a human osteopontin ELISA (R&D Systems). Each dot represents an individual sample. Results are shown as means ± SEM, n=2-20. *p<0.05, **p<0.01; One-Way ANOVA with post-hoc Tukey’s test

In alignment with previous literature,^29^ milk from mothers who gave birth to preterm infants trended toward higher levels of osteopontin than milk from mothers delivering term infants (Figure 2). However, the preterm samples had a greater reduction in osteopontin concentration post-pasteurization (Figure 3).

**Figure 2:**
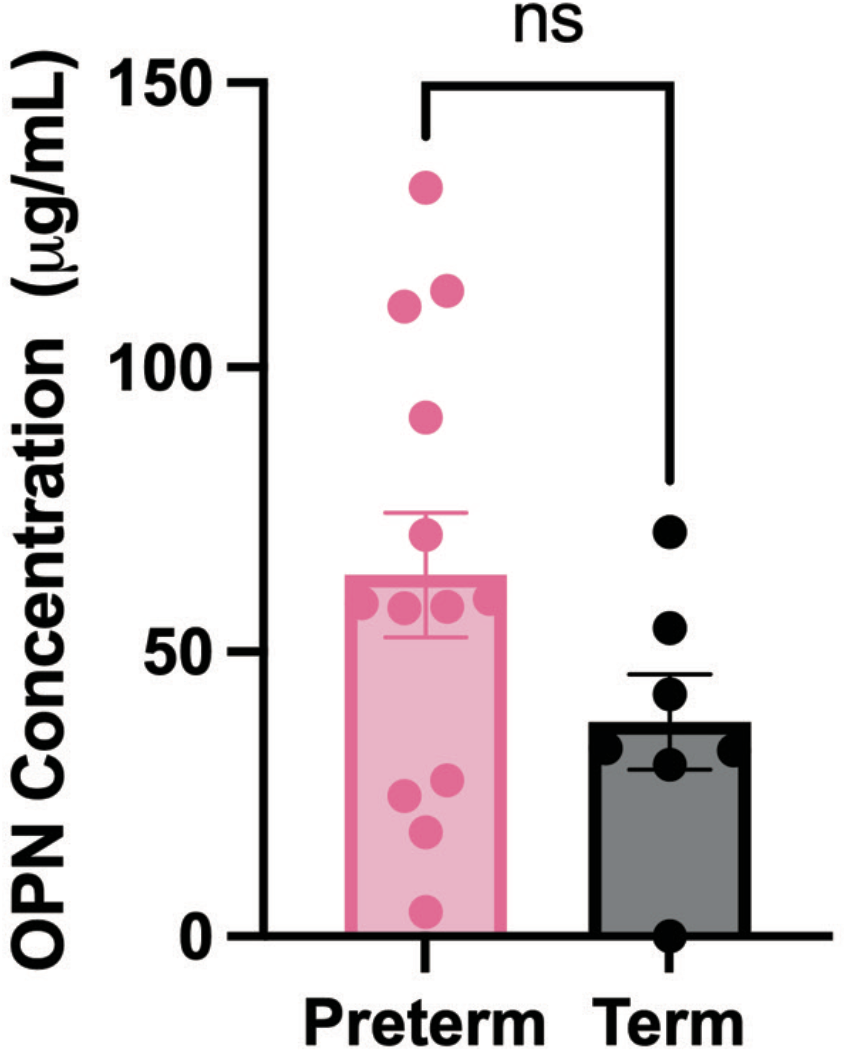
Osteopontin levels in breastmilk from mothers of preterm and term infants. Human milk osteopontin concentrations were measured using a human osteopontin ELISA (R&D Systems). Each dot represents an individual sample. Results are shown as means ± SEM, n=13 preterm, n=7 term.

**Figure 3:**
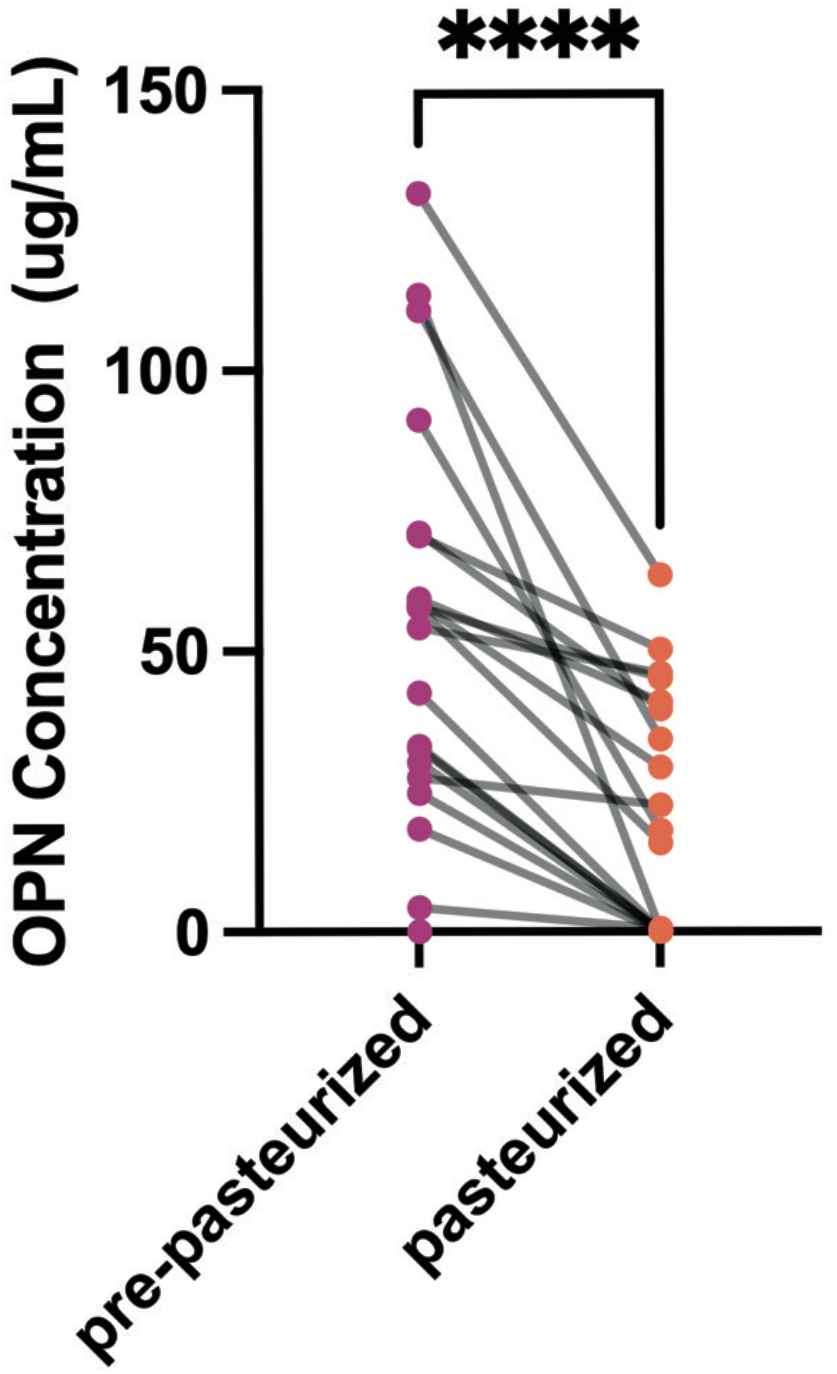
Osteopontin levels in single-donor breastmilk from mothers of preterm and term infants, before and after Holder pasteurization. Samples were pasteurized for 30 minutes at 62.5°C. Human milk osteopontin concentrations were measured using a human osteopontin ELISA (R&D Systems). Each dot represents an individual sample. Results are shown as means ± SEM, n=20. ****p<0.001; Student’s paired t-test.

Freezing and pasteurization are procedures commonly used to preserve milk and ensure safety; however, how the combination of these procedures affects the levels of osteopontin in breastmilk is unknown. To address this issue, a single fresh breastmilk sample was acquired and subjected to combinations of freezing and Holder pasteurization (Figure 4a). Freezing or Holder pasteurization alone resulted in a small decrease in detectable osteopontin, while freezing and pasteurization in combination decreased osteopontin levels by 35-58%. Pasteurization prior to freezing resulted in the lowest osteopontin levels (Figure 4b).

**Figure 4:**
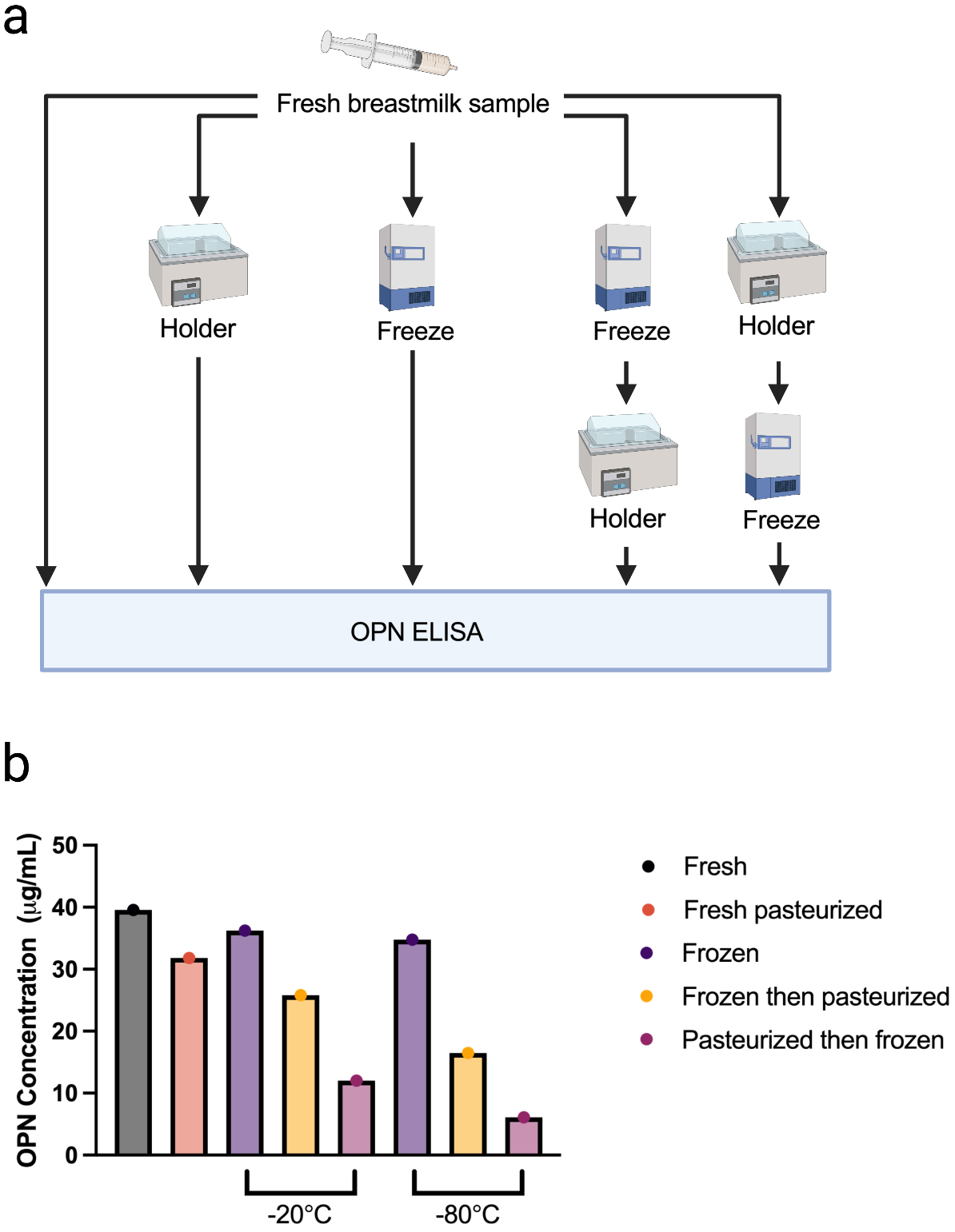
Osteopontin levels in fresh and processed breastmilk. (a) Schematic of breastmilk processing. Breastmilk was aliquoted and subjected to combinations of freezing and Holder pasteurization prior to analysis. (b) Osteopontin concentration in fresh and processed breastmilk.

## Discussion

Because of varying circumstances, not all mothers are able to feed breastmilk to their infants, which has led to use of alternative nutritional sources. Formula has been a common substitute for breastmilk; however, infant formula lacks many bioactive molecules, including osteopontin. Because of these limitations, use of alternate breastmilk products besides MoM has been on the rise. However, it is of high priority to determine whether alternate breastmilk products confer equal nutritional and bioactive value as MoM. Though not recommended by the medical community, single-donor breastmilk is often acquired through personal connections, with a 2018 study demonstrating that approximately 7% of US infants receive donated milk.^30^ In these cases, the Academy of Breastfeeding Medicine advises caretakers to discuss pasteurizing donor milk before consumption.^31^ However, our data and and others^28^ have shown that pasteurization significantly decreases the levels of osteopontin present in milk, which may compromise the biological activities of this protein.

Although single donor breastmilk may be easier to acquire than pooled donor breastmilk, the levels of bioactive molecules present in an individual milk source may not be adequate for infant health, especially after pasteurization (Figure 1). The combination of breastmilk from different donors provides a better alternative to single-donor pasteurized milk because the variance in osteopontin levels among diverse donors “balances” the osteopontin concentration even after Holder pasteurization.

To preserve donor milk for infant consumption, milk is frozen and/or pasteurized. However, our data indicate that the order of these procedures is critical for maintaining “normal” levels of osteopontin. Here we present initial evidence indicating that pasteurization followed by freezing at −20°C (the temperature of most household freezers) significantly depletes osteopontin levels. According to our data, a better alternative is to freeze milk prior to pasteurization, a strategy followed by most milk banks, but which may not be followed by parents at home.

A significant limitation of this study is the lack of precise data about time of milk collection. Due to the deidentified status of the samples, little data was provided regarding at which postpartum stage milk was collected. Thus, we cannot be confident that comparisons between single donor groups are not an artifact of milk collected at different post-partum stages. We were also limited in our acquisition of fresh donor breastmilk samples, which hindered the analysis of osteopontin levels between fresh and frozen samples.

Our data shows that milk pasteurization significantly impacts the concentrations of osteopontin, which may hinder the activity of this biofactor. The functions of milk-derived osteopontin are not clearly established, but the high levels of this protein in breastmilk clearly suggest a pivotal role for this molecule in infant development. Other groups have shown that osteopontin is important for the development of intestinal epithelial cells,^32^ while we have reported that osteopontin is needed for intraepithelial lymphocyte homeostasis.^33^ In addition, it is well known that osteopontin interacts with bacteria,^34,35,36^ which may indicate a role in the establishment of the nascent intestinal microbiota. Taken together, it is possible that milk-derived osteopontin is critical for the development and maturation of intestinal epithelial cells and the mucosal immune system.

Considering these potential functions, it is imperative to consider how donor breastmilk is treated. If freezing and pasteurization are essential for donor milk safety, then osteopontin supplementation might be considered as a potential remediation. In Europe, bovine osteopontin (known as Lacprodan OPN-10), has been approved for formula supplementation, indicating that this protein is well tolerated by infants. Therefore, preterm and term infants that use frozen/pasteurized breastmilk as their main nutritional source may benefit from osteopontin supplementation.

## Data Availability

The datasets generated during and/or analyzed during the current study are available from the corresponding author on reasonable request.

## Acknowledgements

We thank Josh McCrary (Monroe Carell Jr. Children’s Hospital at Vanderbilt) for providing pooled donor breastmilk and retort-pasteurized breastmilk samples. Cartoons in Figure 4 were created with BioRender.com.

## Author contributions

K.M., J.R., and J.W., conceived the project idea; J.W. provided samples; K.M. performed data acquisition and analysis; K.M. and D.O-V. wrote the manuscript; J.R., J.W., K.M., and D. O-V. revised and approved the final article.

## Consent Statement

All samples were acquired from patients who had given informed consent for their samples to be used for research.

